# Evolutionary and structural analyses of SARS-CoV-2 D614G spike protein mutation now documented worldwide

**DOI:** 10.1101/2020.06.08.140459

**Authors:** Sandra Isabel, Lucía Graña-Miraglia, Jahir M. Gutierrez, Cedoljub Bundalovic-Torma, Helen E. Groves, Marc R. Isabel, AliReza Eshaghi, Samir N. Patel, Jonathan B. Gubbay, Tomi Poutanen, David S. Guttman, Susan M. Poutanen

**Affiliations:** Department of Paediatrics, The Hospital for Sick Children, University of Toronto, Toronto, Ontario, Canada; Centre for the Analysis of Genome Evolution & Function, University of Toronto, Toronto, Ontario, Canada; Department of Cell & Systems Biology, University of Toronto, Toronto, Ontario, Canada; Layer 6 AI, Ontario, Canada; Département de mathématique, Université Laval, Québec City, Canada; Public Health Ontario, Toronto, Ontario, Canada; Department of Laboratory Medicine and Pathobiology, University of Toronto, Ontario, Canada; Department of Medicine, University of Toronto, Toronto, Ontario, Canada; Department of Microbiology, University Health System/Sinai Health, Toronto, Ontario, Canada

## Abstract

The COVID-19 pandemic, caused by the Severe Acute Respiratory Syndrome Coronavirus 2 (SARS-CoV-2), was declared on March 11, 2020 by the World Health Organization. As of the 31^st^ of May, 2020, there have been more than 6 million COVID-19 cases diagnosed worldwide and over 370,000 deaths, according to Johns Hopkins. Thousands of SARS-CoV-2 strains have been sequenced to date, providing a valuable opportunity to investigate the evolution of the virus on a global scale. We performed a phylogenetic analysis of over 1,225 SARS-CoV-2 genomes spanning from late December 2019 to mid-March 2020. We identified a missense mutation, D614G, in the spike protein of SARS-CoV-2, which has emerged as a predominant clade in Europe (954 of 1,449 (66%) sequences) and is spreading worldwide (1,237 of 2,795 (44%) sequences). Molecular dating analysis estimated the emergence of this clade around mid-to-late January (10 - 25 January) 2020. We also applied structural bioinformatics to assess D614G potential impact on the virulence and epidemiology of SARS-CoV-2. *In silico* analyses on the spike protein structure suggests that the mutation is most likely neutral to protein function as it relates to its interaction with the human ACE2 receptor. The lack of clinical metadata available prevented our investigation of association between viral clade and disease severity phenotype. Future work that can leverage clinical outcome data with both viral and human genomic diversity is needed to monitor the pandemic.

---

In late December 2019, a cluster of atypical pneumonia cases was reported and epidemiologically linked to a wholesale seafood market in Wuhan, Hubei Province, China^1^. The causative agent was identified as a novel RNA virus of the family *Coronaviridae* and was subsequently designated SARS-CoV-2 owing to its high overall nucleotide similarity to SARS-CoV, which was responsible for previous outbreaks of severe acute respiratory syndrome in humans between 2002-2004^2,3^. Previous studies of SARS-CoV-2 genomes sequenced during the early months of the epidemic (late December 2019 up to early February 2020) estimated the time of its emergence at the end of November (18^th^ and 25^th^)^4–6^, approximately a month before the first confirmed cases. It has been hypothesized that SARS-CoV-2 may have undergone a period of cryptic transmission in asymptomatic or mildly symptomatic individuals, or in unidentified pneumonia cases prior to the cluster reported in Wuhan in late December 2019^3^. Based on the high nucleotide identity of SARS-CoV-2 to a bat coronavirus isolate (96%)^7^, a possible scenario is that SARS-CoV-2 had undergone a period of adaptation in an as yet identified animal host, facilitating its capacity to jump species boundaries and infect humans^3^. The present rapid spread of the virus worldwide, coupled with its associated mortality, raises an important concern of its further potential to adapt to more highly transmissible or virulent forms.

The availability of SARS-CoV-2 genomic sequences concurrent with the present outbreak provides a valuable resource for improving our understanding of viral evolution across location and time. We performed an initial phylogenetic analysis of 749 SARS-CoV-2 genome sequences from late-December 2019 to March 13, 2020 (based on publicly available genome sequences on GISAID) and noted 152 SARS-CoV-2 sequences initially isolated in Europe beginning in February, 2020, which appear to have emerged as a distinct phylogenetic clade. Upon further investigation, we found that these strains are distinguished by a derived missense mutation in the spike protein (S-protein) encoding gene, resulting in an amino acid change from an aspartate to a glycine residue at position 614 (D614G). At the time of our study, the D614G mutation was at particularly high frequency 20/23 (87%) among Italian SARS-CoV-2 sequenced specimens, which was then emerging as the most severely affected country outside of China, with an overall case fatality rate of 7.2%^8^.

During the course of our analysis, the number of available SARS-CoV-2 genomes increased substantially. We subsequently analyzed a total of 2,795 genome sequences of SARS-CoV-2 (Supplementary data Table 1). For those sequences with demographics (65% of sequenced specimens), the male to female ratio was 0.56:0.44, with a median age of 49 years old, and a range from less than 1 to 99 years old. As of 30 March 2020, the D614G clade includes 954 of 1,449 (66%) of European specimens and 1,237 of 2,795 (44%) worldwide sequenced specimens. A comparison against the previous set of genomes collected for our phylogenetic and molecular dating analysis revealed that for samples submitted during the period from March 17-30, 2020, the D614G clade became increasingly prevalent worldwide, expanding from 22 to 42 countries (Figure 1), as also reported previously^9,10^. The demographic distribution for this mutation, when known, (male to female ratio, 0.56:0.44; median age, 48 years old; age range, less than 1 to 99 years old) was not significantly different compared to the reported demographics for all sequenced SARS-CoV-2 specimens.

**Figure 1.**
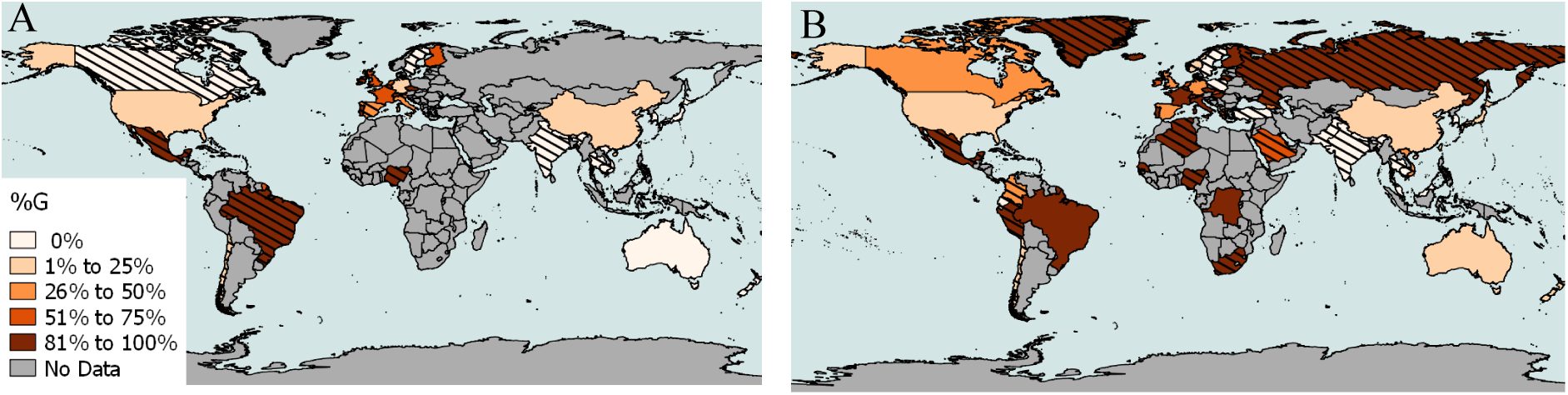
Global Distribution of SARS-CoV-2 Genome Sequences Possessing the Spike Protein D614G Mutation. G mutation as percentage of total sequences (% G) is represented with color shades as detailed in legend including data available as of A) 17 March and B) 30 March 2020. Hatched lines were added when less than 10 sequences were available for one country.

We employed molecular dating to estimate the time of emergence of the D614G clade. Based on a curated set of 442 genomes representing the sequence diversity of SARS-CoV-2 samples available at the time of analysis (30 March 2020), the mean time to most recent common ancestor (tMRCA) was estimated to be on 18 January 2020 (95% highest posterior density (HPD) interval: 10 January - 25 January), indicating its relatively recent emergence (Figure 2). Although the mutation appears to be clade-specific, we noted that D614G also arose another time in a single isolate belonging to a distinct lineage, Wuhan/HBCDC-HB-06/2020 (EPI 412982), collected on 7 February 2020. Given the high degree of nucleotide identity of the D614G clade (∼99.6%), we expect that future tMRCA estimates will not differ substantially. The mean tMRCA for the root of the tree was estimated to be 13 November 2019 (95% HPD interval: 19 October 2019 to 5 December 2019) which is similar to what has been estimated previously^4^. Multiple factors can influence these estimates as more data become available: increased diversity and number of strains included for analysis, expanded range of sampling dates compared to earlier studies, as well as incomplete/biased sampling of available genomes. Therefore, caution should be used when inferring broad and definite conclusions about the epidemiology of the emerging outbreak in real-time, particularly as the early undocumented stages of SARS-CoV-2 transmission is largely unknown^3^. They should be taken as tentative estimates based on limited information, which are subject to change as more information becomes available and compared to other epidemiological information. For instance, as different countries review and test archived specimens from cases of severe pneumonia or influenza-like illness for SARS-CoV-2, it is expected that additional cases may be identified, such as in France where a patient without travel history to China was identified to have COVID-19 in late December concurrent with the initial reported cases from Wuhan^11^. These retrospective analyses will provide crucial insights into the early transmission dynamics and evolution of SARS-CoV-2 and its rapid global spread.

**Figure 2.**
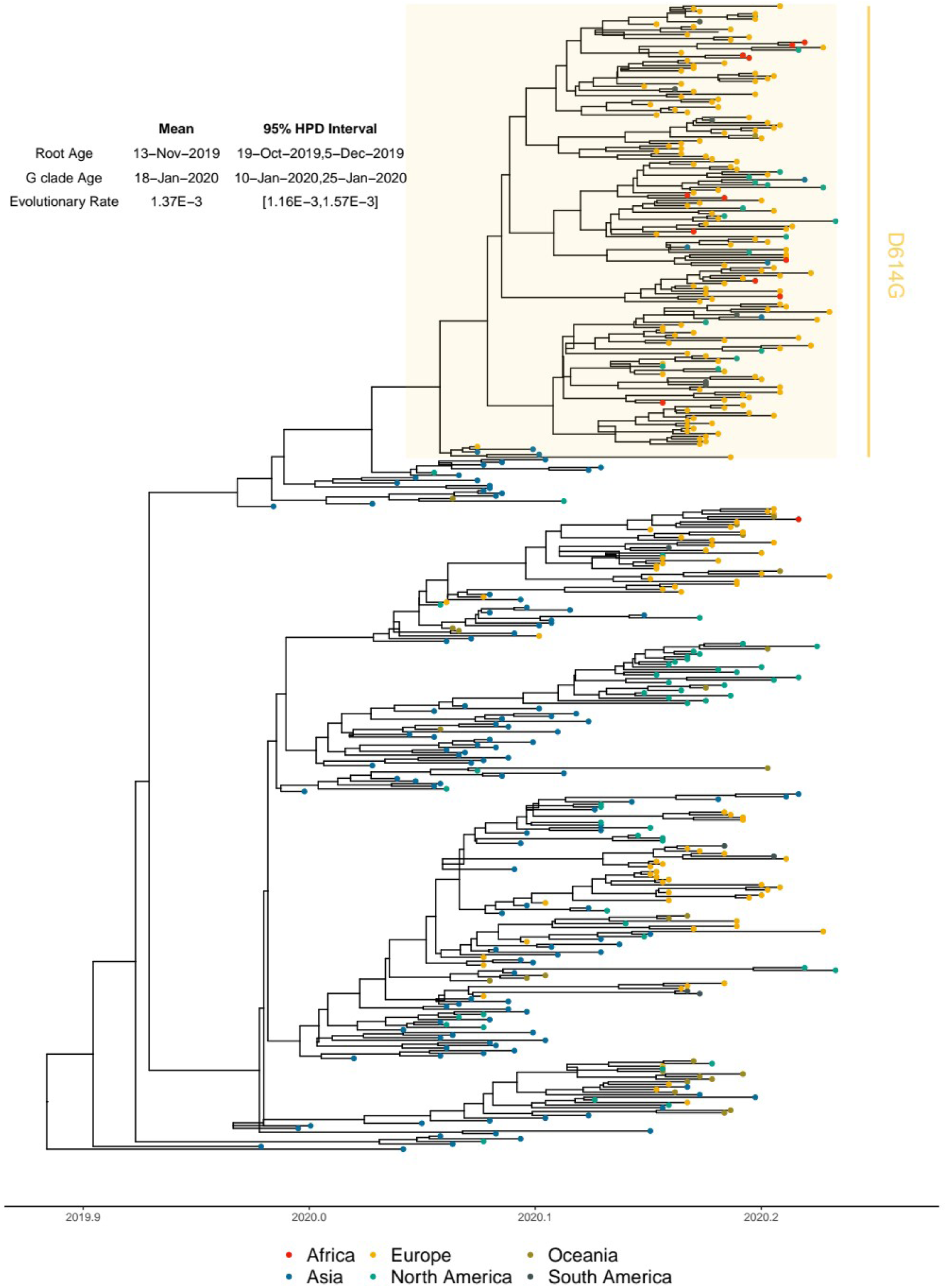
Estimated Molecular Dating of Evolutionary history of 442 Representative Global SARS-CoV-2 Sequences (Late-December 2019 – Mid-March 2020) and the Emergence of the D614G Clade. Maximum clade credibility (MCC) tree with dated branches estimated by Bayesian Evolutionary Analysis Sampling Trees (BEAST). Node colors indicate continents of isolation; x-axis indicating dates by year and days in decimal notation; D614G clade sequences are highlighted in a yellow box.

From our findings of the recent emergence of the D614G clade and the increasing number of specimens harboring the mutation identified worldwide, we sought to investigate the potential significance of the mutation on clinical disease severity phenotypes. Unfortunately, only limited clinical disease severity data was available for patients whose sequenced samples were included in our analysis (asymptomatic 64, symptomatic 8, mild symptoms 2, pneumonia 4, hospitalized 163, released 48, recovered 26, nursing home 3, live 24, ICU 2, deceased 1), preventing us from being able to meaningfully correlate disease severity and genotype from these data. However, using country-wide crude case fatality data for countries from which sequencing data was available, there was no significant correlation between proportion of D614G clade sequences and crude case fatality rate as of 30 March 2020 (Spearman’s rank correlation coefficient, r 0.22, 95% CI -0.12 to 0.51). In addition, on analysis of crude case fatality rate by age-group (available for China, Italy, South Korea, Spain, and Canada) there was no significant correlation with proportion of D614G clades in the sequences analysed for these countries (Supplementary Table 2)^8,12–14^ A major limitation of this analysis is that publicly available SARS-CoV-2 genomes are the result of convenience sampling and are not expected to provide an accurate representation of the spatial-demographic distribution of SARS-CoV-2 genotypes. Therefore, one must be cautious about making inferences about severity and transmission of variants^15^ from genomic sequence data alone.

The possibility that the D614G mutation may still have a potential impact on the function of the SARS-CoV-2 spike protein could not be excluded. Previous studies of SARS-CoV have shown that the sequential accumulation of mutations in the spike protein increased its affinity to ACE2 and likely impacted its transmission and disease severity during the course of outbreaks in 2002-2004^16–18^. Recently, Phan *et al.* studied 86 complete SARS-CoV-2 genomes, identifying 42 missense mutations, eight of which occurred in the S-protein gene^2^. However, at the time of their study, the mutation D614G was only found in one sequence from Germany (collected on 28 January, 2020). Modifications in the spike protein are of interest as they might indicate the emergence of a novel strain of SARS-CoV-2 with change in transmissibility or pathogenicity. Therefore, we investigated the potential functional and epidemiological consequences of the D614G mutation with structural modeling of the SARS-CoV-2 spike (S) protein and its interaction with the angiotensin-converting enzyme 2 (ACE2) receptor.

The structure of the SARS-CoV-2 spike (S) protein is shown in Figure 3. The S protein is a heavily glycosylated trimeric protein that mediates entry to host cells via fusion with ACE2. Recently, Wrapp and colleagues used Cryo-EM to determine the structure of the S protein and analyze its conformational changes during infection^19^. Using their three-dimensional model of the S protein structure, we set out to investigate the effects that a mutation in position 614 might have. First, we analyzed changes in inter-atomic contacts within a radius of 6 Å from position 614 before and after substitution of aspartate by glycine. Notably, four inter-chain destabilizing (i.e., hydrophobic-hydrophilic) contacts are lost with residues of an adjacent chain upon D614G mutation (see Table 1 and Figure 3C). This suggests that a small repelling interaction between adjacent chains is removed upon this aspartate substitution (see Table 1). However, it is unlikely that this would have a significant effect on recognition and binding to ACE2 given the relative distal position of this mutation with respect to the receptor-binding domain (RBD) (see Figure 3B), but further analyses would be required to assess whether the D614G mutation has an effect on the way the S protein changes its conformation after interaction with ACE2. Lan and colleagues also showed residues in the RBD act as epitopes for SARS-CoV-2 and mutations can influence antibody binding^20^. Given the important role that glycosylation plays in regulating the function of spike proteins in coronaviruses^21^, we decided to search for potential changes in a glycosylated residue (asparagine) in position 616. As shown in Figure 3D, this residue and aspartate 614 are oriented in opposite directions, suggesting that substitution by glycine also has a null effect on this interaction. Finally, neither position 614 nor the inter-atomic contacts at positions 854, 859, 860, 861 of the spike protein lie in a polybasic cleavage region which is of importance for SARS-CoV-2 as it has been proposed to activate the protein for membrane fusion^22^. Our results are consistent with the work by Wrapp and colleagues, where they identified nine missense mutations (including D614G) in the spike protein that were thought to be relatively conservative and thus unlikely to affect protein function^19^.

**Table 1.** Inter-chain contacts lost upon D614G mutation between adjacent chains in the SARS-CoV-2 Spike protein.

| Residue in non-reference adjacent chain | Distance (Å) | Contact surface area (Å^2) |
| --- | --- | --- |
| Lys 854 | 5.2 | 10.0 |
| Thr 859 | 2.7 | 28.8 |
| Val 860 | 4.5 | 5.6 |
| Leu 861 | 5.6 | 1.0 |

**Figure 3.**
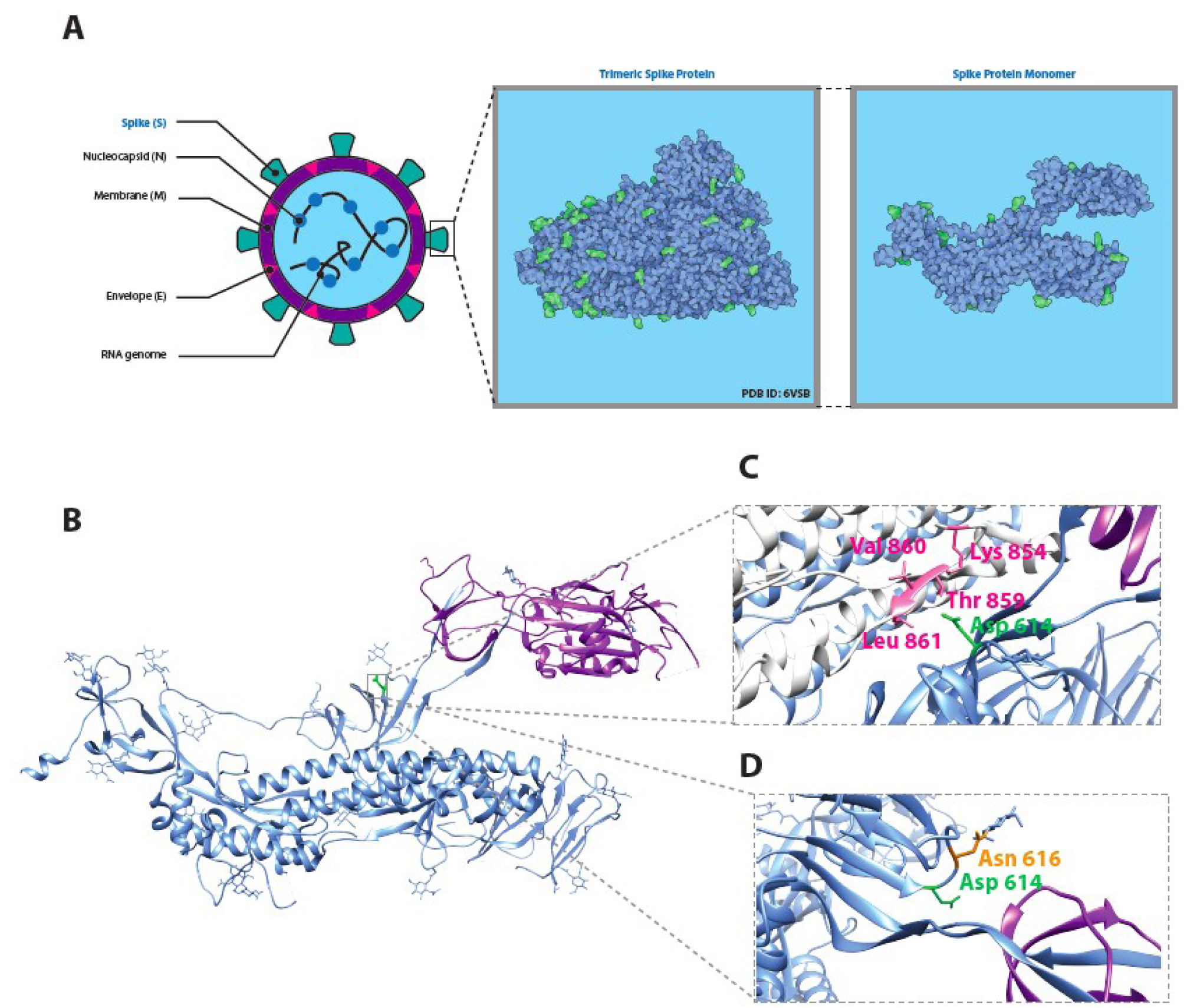
Structural analysis of SARS-CoV-2 spike protein around position 614. (A) Location and distribution of SARS-CoV-2 viral proteins. The full trimeric form of the spike protein results from a complex of three identical spike monomers (right panel). (B) Three-dimensional depiction of a spike protein monomer. The receptor-binding domain is colored purple and the location of the aspartate residue in position 614 is highlighted in green. (C) Inter-atomic contacts between aspartate 614 (green) in a reference spike monomer (blue) and four residues (pink) in its adjacent spike protein monomer chain (white). These four contacts are destabilizing and create a hydrophilic-hydrophobic repelling effect that is lost upon replacement of aspartate by glycine in the D614G mutation (see Table 1). (D) Spatial distribution of aspartate 614 residue (green) and an adjacent glycosylated asparagine residue in position 616 (orange). The two residues point in opposite directions and thus it is unlikely they share a meaningful interaction.

There were important limitations faced in our present analysis which are likely to be a significant hurdle to similar studies in the future. The lack of available clinical metadata prevented our investigation of association between viral clade and disease severity phenotype. Additionally, numbers of sequenced SARS-CoV-2 samples vary greatly between countries and may be subject to potential sampling bias. It is important to note that current country level data on crude case fatality rates and case numbers do not permit robust comparison of clinical phenotype across countries due to significant differences in population demographics, testing protocols, case definitions and implementation of public health measures. Accordingly, it was not possible to draw any conclusions regarding the clinical phenotype of the D614G clade and there is no evidence at this time to suggest this clade is associated with any differences in disease phenotype. As a potential proxy measure for viral load, we reviewed the cycle threshold (Ct) values of the SARS-CoV-2 assay for samples sequenced from Ontario, Canada (total n = 178, with 50 and 128 samples possessing the spike protein 614-D and 614-G mutations, respectively). In this small selection of samples, we found no significant difference in the average Ct for 614-D and 614-G mutations (mean 22.35, SD 5.13 and mean 22.32, SD 5.188 respectively, t-test p-value = 0.978) (Figure 4). We do note, however, that numerous factors affect the measurement and limit the reliability of Ct value measurement, including sample type, quality of swabs, quality of collection method, and time from onset to sample collection^23^. In contrast, another study has found a statistically significant decrease in PCR Ct values associated with the mutated amino acid G variant^10^ but its clinical significance remains unclear. In spite of the lack of evidence for phenotypic differences, it is interesting that in a short period of time since its emergence the D614G clade has become widespread all around the world. It is possible that even if the observed mutation does not impact the protein’s interaction with ACE2, it may not be completely neutral with respect to viral fitness. For example, given that the molecular weight of glycine is significantly smaller than that of aspartate, the mutation could be advantageous from a cost minimization point of view^24^. It is also possible that the mutation may be in linkage disequilibrium with a selectively advantageous variant impacting another aspect of viral reproduction. For example, most strains of the D614G clade also harbor a mutation (P4715L) in orf1ab and a subclade processes three nucleotide changes in the nucleoprotein (N) gene (GGG to AAC; G204R), which plays diverse roles in virion assembly as well as genome transcription and translation^25,26^. Most recently, it has been suggested that D614G introduces a new elastase cleavage site that may be differentially activated by host genomic mutations thereby facilitating spike processing, and entry into host cells in some host populations but not others^27^. Finally, the global spread of the D614G mutation may have nothing to do with viral biology but may simply be a consequence of the high level of interconnectedness of Europe to the rest of the world. Thus, the emergence of the D614G clade may be explained by a founder event and subsequent clonal expansion in Europe that led to its spreading worldwide.

**Figure 4.**
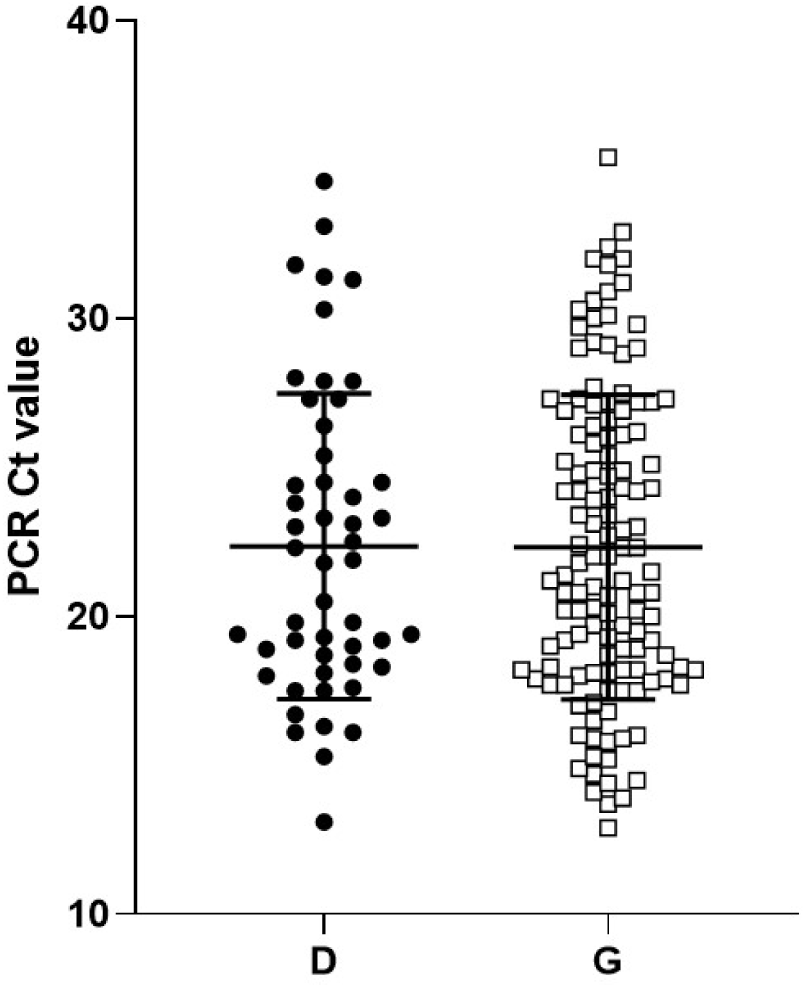
SARS-CoV-2 PCR Cycle threshold (Ct) values of different clinical samples plotted according to variant D (black dots) and G (white squares) at the position 614 in the spike protein. Dots represent individual Ct values; horizontal lines represent the mean and standard deviation.

Genomic sequencing and phylogenetic analysis are powerful approaches to track viral evolution during the course of a pandemic and help coordinate the global implementation of strategies for decreasing virus transmission and mitigating global mortality. However, our study highlights that caution is warranted in hastily drawing conclusions from limited observational data. Future work that can leverage clinical outcome data with both viral and human genomic diversity, *in vitro* and *in vivo* tests and *in silico* modeling will greatly enhance our ability to understand if specific SARS-CoV-2 clades are evolving with respect to virulence or transmissibility, and identify genetic variants associated with viral adaptation, transmissibility, and mortality. These data will be essential for effective vaccine development and public health responses to this emergent pandemic.

## Materials and Methods

### Sequence data, demographics, maps and statistics

A total of 2,803 genomes of SARS-CoV-2 (excluding non-human sequences specimens) and demographics when available were obtained from GISAID (http://gisaid.org/) (downloaded on 30 March 2020) to perform the statistical analyses of the D614G mutation (8 sequences were excluded due to ambiguous or unknown nucleotide at this position). The dataset included sequences representing 55 countries. The world geographical maps were built with the geographic information system QGIS (v2.18.21, https://qgis.org). Global CFR was calculated from data according to Johns Hopkins as of 30 March 2020. CFR for age groups of different countries were extracted from country-specific published data ^8,12–14^. Spearman’s rank correlation analysis was performed using GraphPad prism (v 8.3.1).

### PCR and sequencing

SARS-CoV-2 E-gene PCR on respiratory specimens (e.g. nasopharyngeal swabs and bronchoalveolar lavages) submitted to Public Health Ontario Laboratory was performed as previously described^28^. Specimens with Ct values of 32 or less were selected for whole genome sequencing. Briefly, rRNA depleted RNA was reverse transcribed with SuperScript III Reverse Transcriptase (Invitrogen/Life Technologies, Carlsbad, CA) to cDNA using tagged random primer (5’-GTTTCCCAGTCACGATA- (N9)-3’) followed by second strand DNA synthesis using Sequenase (Thermo Fisher Scientific, Waltham, MA). Amplification of double stranded cDNA was done with primer 5’-GTTTCCCAGTCACGATA-3’ using AccuTaq™ LA DNA Polymerase according to manufacture recommendation. Amplified products were purified using 1.0 X ratio of Agencourt AMPure XP beads (Beckman Coulter) and quantified with Qubit 2.0 Fluorometer by using Qubit dsDNA HS Assay Kit (Thermo Fisher Scientific). Nextera XT DNA Library Preparation kit (Illumina) following manufacturer’s instructions was used for Library preparation and indexing. The sequencing library was quantified using Qubit 2.0 (Invitrogen, Waltham, MA, USA) and average size was determined by 4200 TapeStation using the D5000 ScreenTape assay (Agilent Technologies, US). The library was normalized to 4 nMol and sequenced on the Illumina MiSeq platform, using Illumina MiSeq reagent kit v2 (2 × 150 bp), according to the manufacturer’s instructions. CLC Genomics Workbench version 8.0.1 (CLC bio, Germantown, MD, USA) was used for both reference assembly and de novo assembly. Additionally, we used protocol published by Artic Network^29^. The method is based on overlapping specific primers producing short amplicon covering entire genome. Briefly, cDNA synthesis and amplicon generation using 2 individual pools of primers was done according to the published protocol^29^. The two pools of amplicons were combined together and cleaned by Agencourt AMPure XP beads (Beckman Coulter) prior to library preparation using Nextera XT (Illumina) following the manufacturer protocol except that half volume of the reagent was used throughout the protocol. Agencourt AMPure XP beads (Beckman Coulter) in ratio of 1.0 were used for the final purification step followed by measurement by Qubit 2.0 and average size determination, using 2200 TapeStation (Agilent Technologies, USA). All libraries were manually normalized to 4 nMol. Libraries were combined in equal volumes for denaturation and subsequent dilution according to MiSeq protocol recommended by manufacturer.

### Molecular evolution

A total of 2,803 SARS-CoV-2 genomes were obtained from GISAID (downloaded on March 30th 2020). Initial quality filtering was performed by iteratively removing redundant sequences with 100% nucleotide identity, having less than 2,940 informative positions, and lacking informative sampling dates. The resulting 2,017 curated sequences were aligned using mafft^30^ (v7.450: --retree 2 –maxiterate 2 –auto settings), and unaligned ends and gap positions from the resulting alignment were removed using Geneious (version R10.2.6, https://www.geneious.com). A Maximum-likelihood tree was constructed using IQTree^31^ (v1.6.1: -model HKY) and used for identification and removal of highly divergent sequences with Treetime^32^ (clock setting). Following previous recommendations (adapted from http://virological.org/t/issues-with-sars-cov-2-sequencing-data/473)^33^ to account for regions which might potentially be the result of hypervariability or sequencing artifacts, alignment positions showing significant homoplasy were first identified using Treetime (homoplasy setting). Top-10 significant homoplastic positions were merged from three separate analyses, using all sequences from the initial alignment, and sequence subsets generated using Illumina (1,450 samples) or Nanopore (341 samples) sequencing technologies. Next, recombinant clusters of mutations were identified with ConalFrameML^34^. Identified positions were masked using a custom-written python script, and further filtered for sequences with 100% nucleotide identity, resulting in an alignment of 1,225 sequences. Finally, to reduce sampling bias only one sample per country was kept for date of sampling, leading to 442 sequences. Molecular dating analysis was performed by the Markov chain Monte Carlo (MCMC) method implemented by Bayesian Evolutionary Analysis on Sampling Trees (BEAST) 1.10.4^35^. A HKY85 nucleotide substitution model was used. Fixed and relaxed molecular clock models were fitted, and constant population size was compared with exponential population growth. All models were run using default priors except for the exponential growth rate (Laplace distribution) in which scale was set to 100. The chain length was set to 100 million states and burn-in of 10 million. Convergence was checked with Tracer 1.7.1^36^. Based on ESS values, estimates for tMRCA of the European Clade spike protein mutation and root age are reported under the strict molecular clock and constant population growth (ESSs > 900). The resulting coalescent tree was generated using TreeAnnotator^37^ and visualized using ggtree package^38^ in R version 3.6.1 (https://www.r-project.org/).

### Structural analyses

Visualization, analysis and *in silico* mutations of protein structures were done in UCSF Chimera^18^. First, we downloaded the molecular structure of the spike protein from the Protein Data Bank (PDB). This structure corresponds to that resolved by Wrapp and colleagues^19^ and deposited in PDB with identifier number 6vsb. We replaced the aspartate residue in position 614 by glycine using the “Rotamers” function in Chimera with default parameters. Then, inter-atomic contacts of both residues at position 614 were derived with the CSU package^19^ freely available at http://oca.weizmann.ac.il/oca-bin/lpccsu.

## Acknowledgments

We are very grateful to Dr William P. Hanage for his expert comments and advice on the preparation of this manuscript. We would like to thank Dr James M. Rini for protein function analysis and Manon Tetreault for data collection. We are grateful to the numerous scientists who have shared their genome data with the world on GISAID (Supplementary data Table 3). Funding was provided by the Canadian Institutes of Health Research and the Natural Sciences and Engineering Research Council of Canada Collaborative Health Research Project grant [CP-151952] and Public Health Ontario.

## AUTHORS’ CONTRIBUTIONS

SI, HEG, SMP: study design, data analyses, manuscript redaction

LG-M, CBT, DSG: study design, phylogenetic analyses, manuscript redaction

JBG, TP: study design, protein function analyses, manuscript redaction

MRI: geographic data analysis, manuscript redaction

AE, SNP, JBG: study design, sequencing, manuscript redaction

## FUNDING

Funding was provided by the Canadian Institutes of Health Research and the Natural Sciences and Engineering Research Council of Canada Collaborative Health Research Project grant [CP-151952] and Public Health Ontario.

## DISCLOSURE

### Financial disclosure

Dr. Gubbay has received research grants from GlaxoSmithKline Inc. and Hoffman-La Roche Ltd to study antiviral resistance in influenza, and from Pfizer Inc. to conduct microbiological surveillance of *Streptococcus pneumoniae*. These activities are not relevant to this study. Dr. Poutanen has received honoraria from Merck related to advisory boards and talks, honoraria from Verity, Cipher, and Paladin Labs related to advisory boards, partial conference travel reimbursement from Copan, and research support from Accelerate Diagnostics and bioMerieux, all outside the submitted work. All other authors declared that they have no competing interests.

## Figures and Tables

**Supplementary data Table 1.**
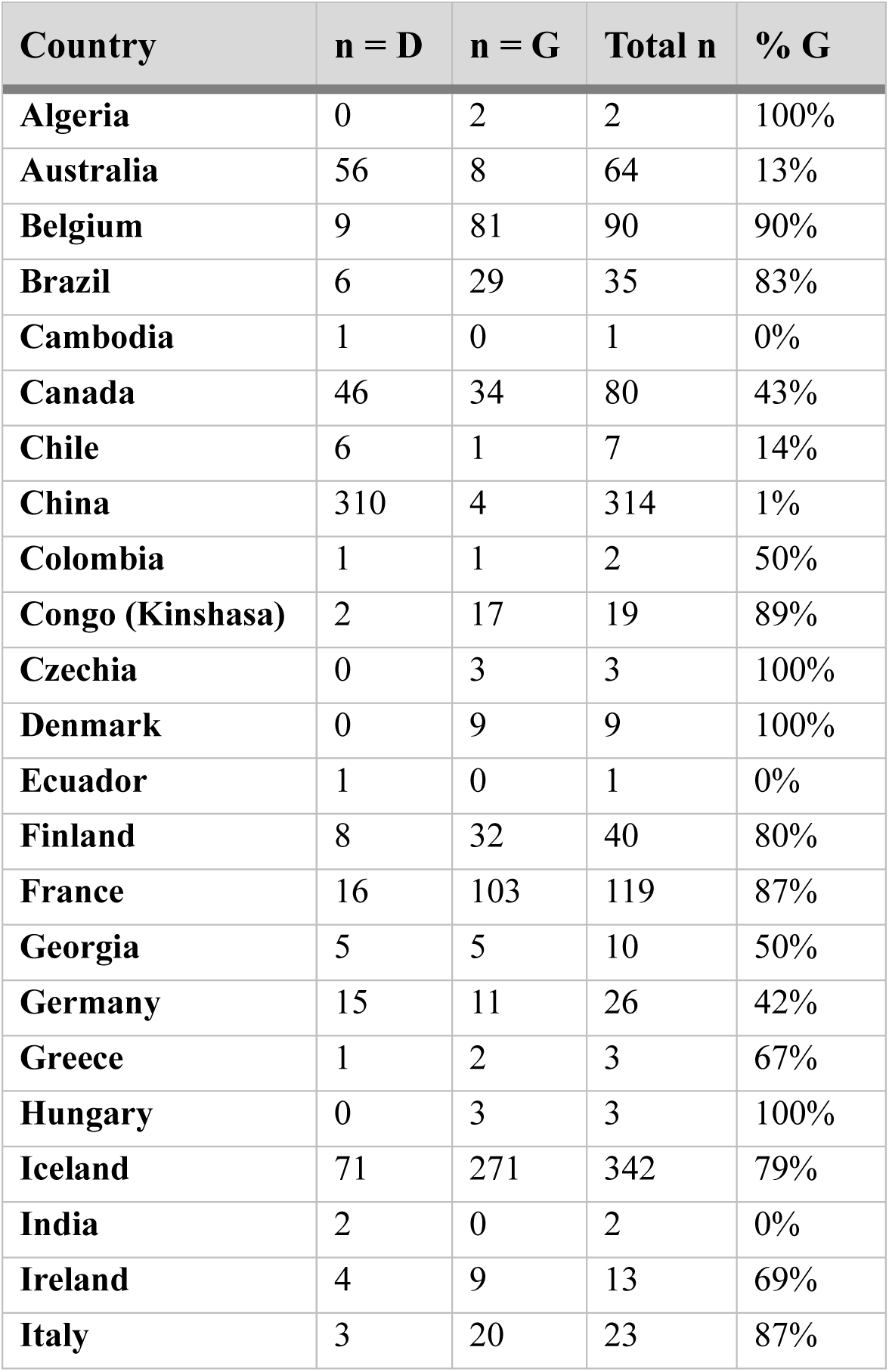
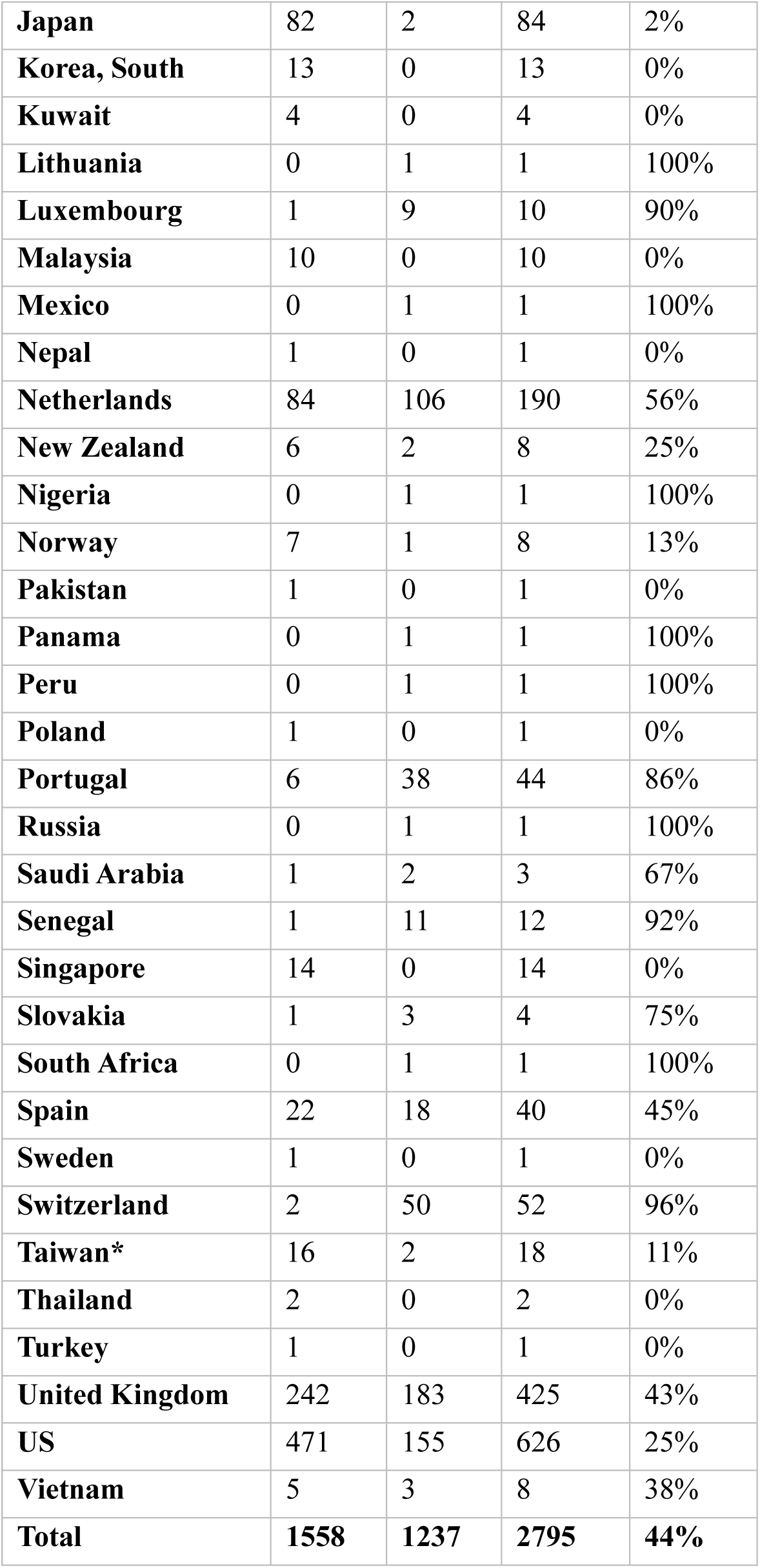
Country Distribution of SARS-CoV-2 Genome Sequences Possessing the Spike Protein D614G Mutation.

**Supplementary data Table 2.** Crude Fatality Rate by age and percent of D614G mutation for China, Italy, South Korea, Spain and Canada.

| Age group (yrs) | Case Fatality Rate by country (%) |  |  |  |  | Spearman's rank correlation r | P-value |
| --- | --- | --- | --- | --- | --- | --- | --- |
|  | China as of 11 Feb 2020 | Italy as of 17 March 2020 | South Korea as of 12 March 2020 | Spain as of 10 April 2020 | Canada as of 9 April 2020* |  |  |
| ≥ 80 | 14.8 | 20.2 | 8.5 | 21.5 | 12.3 | 0.80 | 0.33 |
| 70-79 | 8 | 12.8 | 5.0 | 10.7 | 2.8 | 1.00 | 0.08 |
| 60-69 | 3.6 | 3.5 | 1.5 | 3.4 |  | 0.40 | 0.75 |
| 50-59 | 1.3 | 1 | 0.4 | 1.0 | 0.3 | 0.32 | >0.99 |
| 40-49 | 0.4 | 0.4 | 0.1 | 0.4 |  | 0.77 | 0.50 |
| 30-39 | 0.2 | 0.3 | 0.1 | 0.2 | 0.1 | 0.95 | 0.17 |
| 20-29 | 0.2 | 0 | 0 | 0.2 |  | 0 | >0.99 |
| 10 to 19 | 0.2 | 0 | 0 | 0.2 | 0 | 0 | >0.99 |
| 0-9 | 0 | 0 | 0 | 0.3 |  | 0.26 | >0.99 |
| All ages | 2.3 | 7.2 | 0.9 | 6.5 | 2.3 | 1.00 | 0.08 |
| Percent of G (%) |  |  |  |  |  |  |  |
| All ages | 1.2 | 86.9 | 0 | 45.0 | 42.5 | - | - |

\*Canadian data presented in 20 years ranges were excluded from Pearson correlation analysis.

## Notes

### Competing Interest Statement

The authors have declared no competing interest.

